# ROS accumulation in cotton ovule epidermal cells is necessary for fiber initiation

**DOI:** 10.1101/000745

**Authors:** Mingxiong Pang, Nicholas Sanford, Thea A Wilkins

## Abstract

Cotton (*Gossypium hirsutum*) fiber, an extremely elongated and thickened single cell of the seed epidermis, is the world’s most important natural and economical textile fiber. Unlike *Arabidopsis* leaf trichomes, fiber initials are randomly developed and frequently form in adjacent seed epidermal cells and follow no apparent pattern. Numerous publications suggested cotton fiber development shares a similar mechanism with *Arabidopsis* leaf trichome development. Here we show that H2O2 accumulation in cotton ovule epidermal cells by NBT staining ovules at different development stages between TM1 and N1n2, a lintless-fuzzless doubled mutant originated from TM1. In contrast, *Arabidopsis* and cotton leaf trichomes do not show H2O2 content. By adding DPI (H2O2 inhibitor) and SHAM (H2O2 activator) in vitro ovule cultures, we show fiber initiation directly involves with H2O2 accumulation. We propose that the directional accumulation of H2O2 in cotton ovule epidermal cell is the drive for fiber initiation, elongation.

## Main Text

Seed dispersal is sometimes split into autochory (when dispersal is attained using the plant’s own means) and allochory (when obtained through external means). Wind dispersal (anemochory) is one of the autochories and more primitive means of dispersal and plant seeds have a feathery tissue attached to their seeds and can be dispersed long distances. Primitively, plants distributed the seeds majorly through autochory; as animals emerge and human activity, some plants lost their ability to produce seed secondary tissues helpful for dispersal and some plants gained the production for specific seeds tissues such as cotton seed trichomes as human activity. But we do not understand how these tissues develop. Cotton (*Gossypium spp.)* fibers are unicellular and single-celled seed trichomes of economic importance (*1*) and provide a special single-cell functional genomics research platform (*2*, *3*) for fundamental biological processes such as organogenesis and cell wall development in plants (*4*, *5*). Fiber development initiates on the day of anthesis (0 dpa) and 30-40% of primordial cells (*6*) on the ovule epidermis form cuboidal outgrowths called fiber “initials” (*7*). Unlike *Arabidopsis* leaf trichomes, fiber initials is randomly distributed and follow no apparent pattern as fiber initials frequently form in adjacent epidermal cells (*8*,*9*). Fiber initials rapidly elongate to produce highly tapered cells within 2-3 days of anthesis through a biased diffuse growth mechanism (*10*).

The initiation of single celled cotton cuboidal fiber cells is a biologically important and complex process, but how the fiber initiate and how the genes involved are regulated remain elusive. Zhang et al., (*11*) demonstrated by adding 30% H2O2 to in vitro cultured ovules of XinFLM, a linted-fuzzless mutant, that the fiber initiation is partially rescued, but no evident effect on N1, the dominant naked seed mutant from TM-1 genetic background. This implies the H2O2 biosynthesis pathway may be defect and the signal transduction is still functional in XinFLM. Like cuboidal outgrowth fiber initials, Andriunas et al., (*12*) shows a flavin-containing enzyme (NAPDH oxidase) and superoxide dismutase cooperatively generate a regulatory H2O2 to govern ingrowth wall formation, a transfer cell (*13*), using Visia Faba cotyledon culture. Forman et al., (*14*) verified that ROS accumulates in growing wild type root hairs but dramatically decreases in rhd2 short hair mutants and the generation of ROS is inhibited and mimics rhd2 mutant and further showed (*15*) a RhoGTPase GDP dissociation inhibitor controls spatial ROS production during root hair development.

To determine whether H2O2 is necessary for cotton fiber initiation, we firstly used SEM to observe fiber initiation and development dynamics starting ovules at 2 days of pre-anthesis to 2 days of post-anthesis. In the dominant N1 mutant (Fig. 1), no fiber initials are present on ovule at anthesis and very rare fiber initials occur on the ovules at 2 days of post anthesis. The recessive mutant n2 does not show significant change for fiber initials. The double mutants N1n2 generated by crossing N1 and n2 and selecting till 7 generation does not show any fiber initiation. XZ142-fl, a lintless and fuzzless mutant originated from its wild type XZ142, also does not produce any fibers.

**Fig. 1.**
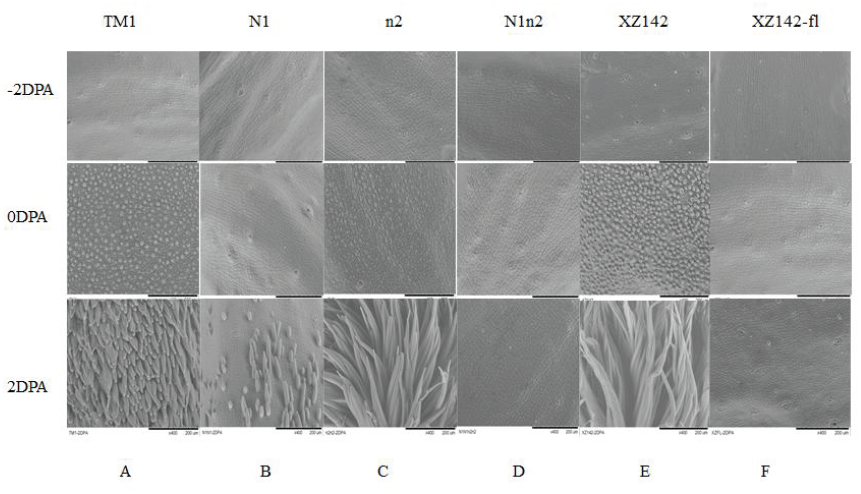
Altered fiber initiation and development in two pair of naked seed mutants with their wild types. (A) Almost all fiber initials occur at 0 DPA in wild type TM1. (B) Few fiber initials occur at 0 and 2 DPA in dominant mutant N1. (C) Many fiber initials occur at 0 and 2 DPA in receive mutant *n2.* (D) No fiber initials developed on *N1n2* double mutant. (E) Almost all fiber initials occur at 0 DPA in wild type XZ142. (F) No fiber initials produced on ovules of XZ142-fl. Black Bar = 200um.

In higher plants ROS can probably be produced by several alternative pathways (*16*) including a cell-wall-localized peroxidase, amine oxidases and non-flavin NADPH oxidases, or NADPH oxidases that resemble the flavin-containing type that is activated by Rac in leukocytes (*17*). Simultaneously, a vast network of antioxidants is acting like ROS scavengers of H2O2 and maintains ROS homeostasis (*18*). This scavenging system consists of catalase (CAT), ascorbate (APX) and secretory peroxidases (POX), glutathione reductases (GR) and peroxiredoxines (PRX), and non-enzymatic compounds like tocopherols, ascorbic acid and flavonoids. (*19*). Li et al., (*20*) identified a cotton cytosolic APX1 (*GhAPX1*) to be highly accumulated during cotton fiber elongation by proteomic analysis and showed (*21*) ethylene is induced in in vitro cultured ovules by endogenous H2O2 and found a feedback regulatory system between H2O2 and ethylene in fiber development. Hovav et al., (*22*) found three genes modulating hydrogen peroxide levels were consistently expressed in domesticated and wild cotton species with long fibers, but not detected by q-PCR in wild species with short fibers. Potika et al., (*23*) Potikha et al., 1999) speculated H2O2 may function as a developmental signal in the differentiation of secondary walls in cotton fibers with support of coincidence of H2O2 generation with the onset of secondary wall deposition and prevention the wall differentiation because of inhibition of H2O2 production or removing the available H2O2. H2O2 modulates downstream signaling events including calcium mobilization protein phosphorylation and gene expression (*24*) which may lead to cotton epidermal cell fate determination. To determine whether H2O2 involves fiber initiation, we used 0.1mg/mL NBT (Nitro Blue Tetrazolium), a chemical specifically staining H2O2 in living cells (Fig.2). Fig.2A timely profiled a normal cotton fiber development starting 5 days of pre-anthesis to fiber maturation at 50 days of post-anthesis. Fig.2B demonstrated that TM1 ovules stained darkly at 2 days of pre-anthesis to 2 days of post-anthesis has already accumulated H2O2 in comparison to these from N1n2. Significantly, in contrast to TM1 ovules, no any staining was detected on *Arabidopsis* and Cotton leaf trichomes (Fig.2C). We also used cut cotton leaf edge as positive, dark staining was found at the cut leaf edge and spots with high concentration of polyphenols (Fig.2C). To rule out the possibility of staining accessibility, we conducted H2O2 production measurement using an Amplex red hydrogen peroxide/peroxidase assay kit (Molecular Probes, http://probes.invitrogen.com/). We found significant decrease of H2O2 in ovules of different stages of N1n2 comparing with those from TM, especially at the 2 DPA (Fig.2D).

**Fig. 2.**
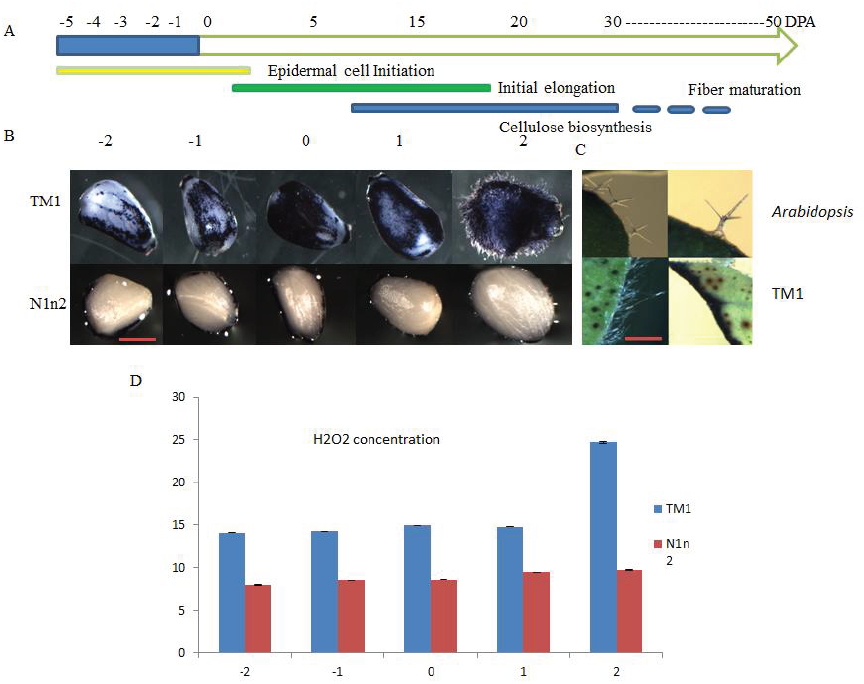
Alteration of H2O2 accumulation by NBT staining and measurement. (A) The time scale indicates the duration of epidermal cell initiation, fiber initial elongation, cellulose biosynthesis and maturation. (B) Ovules from TM1 and Nln2, stained with NBT (C). Leaf trichomes from Arabidopsis and TM1 and TM1 leaf cut edge, stained with NBT. (D). More H2O2 accumulation in ovules from TM1 at different stages comparing to those from Nln2.Red Bar=l mm.

To verify whether H2O2 directly influence epidermal cell differentiation and initiation, we conducted in vitro ovule cultures by adding DPI (H2O2 inhibitor) and SHAM (H2O2 activator. The result (Fig.3A) shows the fiber initiation normally developed in the control (with 0.1% DMSO) in vitro ovule culture and NTB staining shows H2O2 accumulation in the fiber initiation and epidermal cells. Fig.3B shows the fiber initiation was almost totally retarded in the in vitro ovule culture with addition of 0.3 µM DPI and the NTB staining shows barely H2O2 accumulation. Interestingly, Fig.3C shows very similar results with Fig.3A. We cannot conclude SHAM enhances fiber initiation as a more controlled experiment is needed for this result. Based on this data, we can efficiently conclude that H2O2 directly involves cotton fiber initiation and development.

**Fig. 3.**
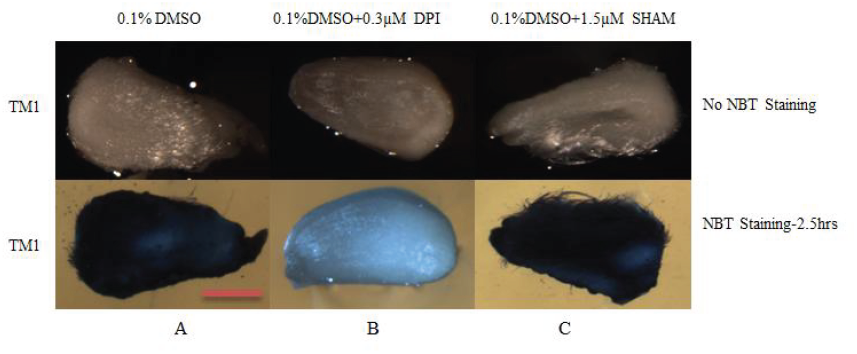
In Vitro Ovule Cultures Treated with DPI and SHAM and Stained with NBT. Cotton ovules from TM1 plants were collected at 0 DPA, treated with DPI (inhibitor at 0.3μM) or SHAM (activator at 1.5 mM) dissolved in DMSO separately for 48 Hrs, and stained with NBT for 2.5 Hrs. Red Bar=l mm.

To molecularly characterize the mechanism of fiber initiation, many researchers (*25*, *26*) proposed the mechanism similarity of cotton fiber development to *Arabidopsis* trichome and study cotton fiber initiation by cloning cotton orthologos for *Arabidopsis* trichome development. Three criteria were used to characterize whether the genes of interest relate to cotton fiber initiation including (i) orthologs to *Arabidopsis* trichome related genes; (ii) highly expressed in cotton ovules and fiber initials and (iii) specific expression in *Arabidopsis* transgenic Col-0 trichome by cotton orthologous promoter-driven GUS and complementation of trichome development for *Arabidopsis* trichomeless mutants by overexpression of cotton ortholog and the varied influences to cotton fiber development in loss-of-function and gain-of function transgenic cotton (*27*). Not like *Arabidopsis* trichome, cotton fibers are initiated from seed epidermal cells, their initiation and development highly synchronize and unbranch with extreme elongation (*28*). To verify the reasonability of the speculation, we roughly grouped the documented cotton fiber development related genes into three groups (S-table 1) including ROS-related, fiber initiation-related and fiber elongation-related.

Transcript profiling for ROS-related genes including *Ghmir398*, with its two targets, *GhCSD1* and *GhCSD2*, *GhAPX*, *GhRDL1* and *GhRDL2*, only *GhRDL1* and *GhRDL2* show consistently lowest expression in the two lintless fuzzless mutants but significantly increase expression starting 2 days of pre-anthisis and post-anthesis (fig.1S). The expression of these genes helps maintain homoestatisis of H2O2, an important signal for cotton fiber initiation and development. We grouped cotton orthologs based on homology to Arabidopsis leaf trichome development into cotton fiber-initiation-related genes. We do not find specific correlation of these group genes to the two lintless fuzzless mutants. Interestingly, all of these genes express in the cotton expanded leaves which produce trichomes (fig.2S). This result suggests these genes may be cotton leaf trichome orthologs but not cotton seed fiber genes.

Interestingly, Walford et al., report (*29*) RNAi cotton transgenic plants for *GhMYB25-like* resulted in fibreless seeds but developed with normal trichomes elsewhere, mimics the Xu142 fl mutant. Furthermore, these plants had decreased expression of the fibers related MYBs including *GhMYB25* and *GhMYB109*, suggesting these *MYBs* are downstream of *GhMYB25-like*. *GhMYB25-like*, not an ortholog to Arabidopsis trichome genes, is identified from its reduced expression in a fibreless mutant of cotton (Xu142 fl) (*30)* Du et al., 2001). These data implicate cotton fiber initiation and development may share similar regulatory process with *Arabidopsis* trichome but the involved genes are different. It is important to note that Arabidopsis seeds do not have seed trichomes like cotton and thus cannot be considered as an ideal surrogate model to evaluate the specificity of expression of cotton fiber genes. The search for cotton fiber initiation and development genes is still elusive. We also detect the expression of the third group of documented fiber related gene, fiber elongation genes, lowest to undetectable expression of *GhMYB25* and *GhEXP1* (fig.3S) relate to the two lintless fuzzless mutants. This implies these two genes are acting downstream and necessary for fiber elongation.

Cell morphogenesis is the process by which cells acquire shape during development and is central to organogenesis in multicellular organisms. Based on the currently available results, Qin et.al. (*31*) speculated that fiber cells may elongate via a combination of both tip-growth and diffuse-growth modes and named linear cell-growth mode. Using NBT staining ovules at different development stages between TM1 and N1n2, a lintless-fuzzless doubled mutant originated from TM, we show that H2O2 accumulation in cotton ovule epidermal cells but no H2O2 staining in *Arabidopsis* and cotton leaf trichomes. We propose that the directional accumulation of H2O2 in cotton ovule epidermal cell is the drive for cotton fiber initiation and elongation.

## Acknowledgments

We thank Dr. Noureddine Abidi for help with Scanning Electron Microscopy (SEM) at Fiber and Biopolymer Research Institute (FBRI) at Texas Tech University and funding by the Texas Governor’s Emerging Technology Superior Research Award granted to T.A.W.

## Supplementary Materials

### Materials and Methods

#### Plant Materials and Scanning Electron Microscopy

Upland cotton (*Gossypium hirsutum L.* cv TM-1), its isogenic naked seed mutants (N1N1 and n2n2), the double mutant N1n2 and XZ142 with it fiberless fuzzless mutant XZ142-fl were grown in the greenhouse under 32/21°C (day/night) temperature regime. Ovules were collected at −2, 0 and 2 dpa flowers at 9 and 10 A.M. Hitachi tabletop TM-1000 scanning electron microscope was used for imaging the fresh ovules.

#### Grouping and QRT-PCR Primers of Potential Genes Related to Fiber Development

We grouped the potential genes related to fiber development into three categories including ROS-related, fiber initiation and elongation (Table-S1).

#### Imaging and ROS Staining for TM1 and N1n2 Ovules at -2DPA to 2DPA

Ovules from TM1 and N1n2 at −2, −1, 0, 1 and 2 DPA were covered with freshly made NBT (nitroblue tetrazolium) solution (in 0.1M potassium phosphate at pH7). Plates were left for 2.5 hrs at room temperature for the color to develop. Ovules were imaged under bright-field illumination using an Olympus SZH10 Zoom Stereo Microscope.

#### H202 Measurement

To measure H2O2 concentration, ovules at different stages were ground in liquid nitrogen, and 200 mg of ground tissues from each sample was added to 200 μL of 20 mM sodium phosphate buffer (pH 6.5) and mixed well and centrifuged at 9500g for 10 min. at 4°C. The supernatants were used for assay. H2O2 concentration was measured using an Amplex red hydrogen peroxide/peroxidase assay kit (Molecular Probes: http://probes.invitrogen.com/).

#### In Vitro Ovule Culture and Treatment with H2O2 Inhibotr and Activator

Cotton ovules from TM1 plants were collected at 0 DPA, soaked in 70% ethanol for 1 min, rinsed in distilled and deionized water and placed in liquid media formulated by Beasley and Ting (*S8*) in a 6-well plate under aseptic conditions. The ovules were cultured in vitro, as described by Meinert and Delmer (*S9*). DPI (inhibitor at 0.3μM) or SHAM (activator at 1.5 mM) dissolved in DMSO was added to medium separately. The final concentration of DMSO was 0.1% in all of the samples.

#### RNA Preparation and QRT-PCR

Total RNA was extracted from ovules, young expanded leaves using the Spectrum™ Plant Total RNA Kit and on-column DNase I digestion, following the manufacturer’s recommendations (Sigma-Aldrich). RNA was quantified using nondrop-1000 and the integrity of total RNA was confirmed by denaturing agarose gel electrophoresis. QRT-PCR amplification reactions were performed using SYBR Green detection chemistry and run in triplicate on 96-wells plates with the 7500 Real Time PCR system (Applied Biosystems). Reactions were prepared in a total volume of 20 μL containing: 5 μL of template, 1 μL of each amplification primer (10uM), 10 μL of 2× Power SYBR Green PCR Master Mix (Applied Biosystems) and 3 μL PCR-quality ddH2O. Cotton Histone-3 (AF024716) was used to normalize the amount of gene-specific RT-PCR products. All PCR reactions were conducted in three replicates and monitored the absence of primer dimers using a dissociation analysis. The amplification data were analyzed using ABI7500 SDS software (version 1.2.2), and the relative expression levels to expression of Histone-3 (fold changes) were calculated. For detecting expression of *GhmiRNA398*, protocol followed (*S1*) using QRT-PCR primer GTCGTATCCATGCAGGGTCCGAGGTATTCGCACTGGATACGACAAGGGG and forward primer ACATTCAAGTGTGTTCTCAGGTCA and universal QRT-PCR miRNA reverse primer GTGCAGGGTCCGAGGT.

**Fig. S1.**
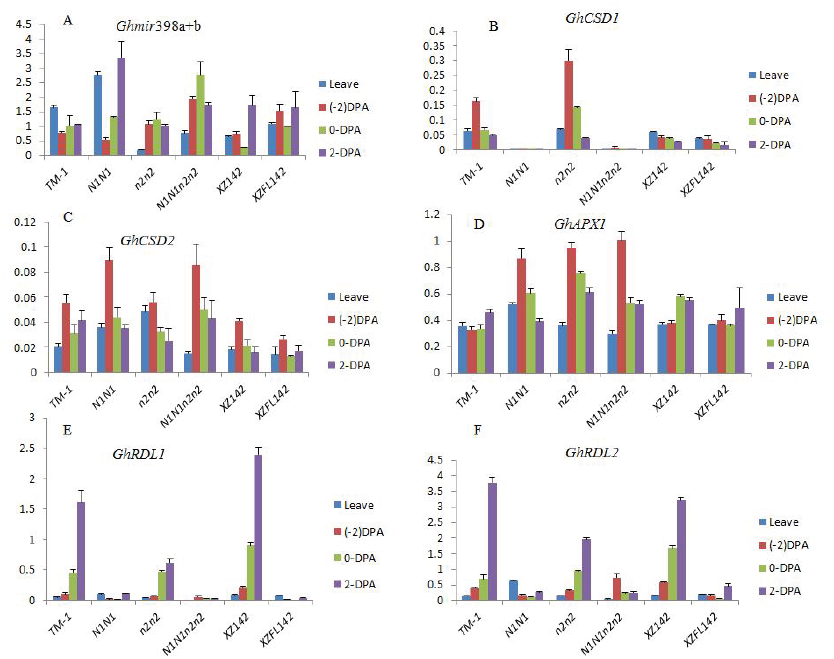
Transcripts profiling for ROS homeostasis-related genes. (A) Gene expression for *Ghmir398a+b* in different varieties and development stages. (B) Gene expression for *GhCSD1(S2)* in different varieties and development stages. (C) Gene expression for *GhCSD2* (*S2*) in different varieties and development stages. (D) Gene expression for *GhAPX1* (*16*) in different varieties and development stages. (E) Gene expression for *GhRDL1* (*26*) in different varieties and development stages. (F) Gene expression for *GhRDL2*(*26*) in different varieties and development stages.

**Fig. S2.**
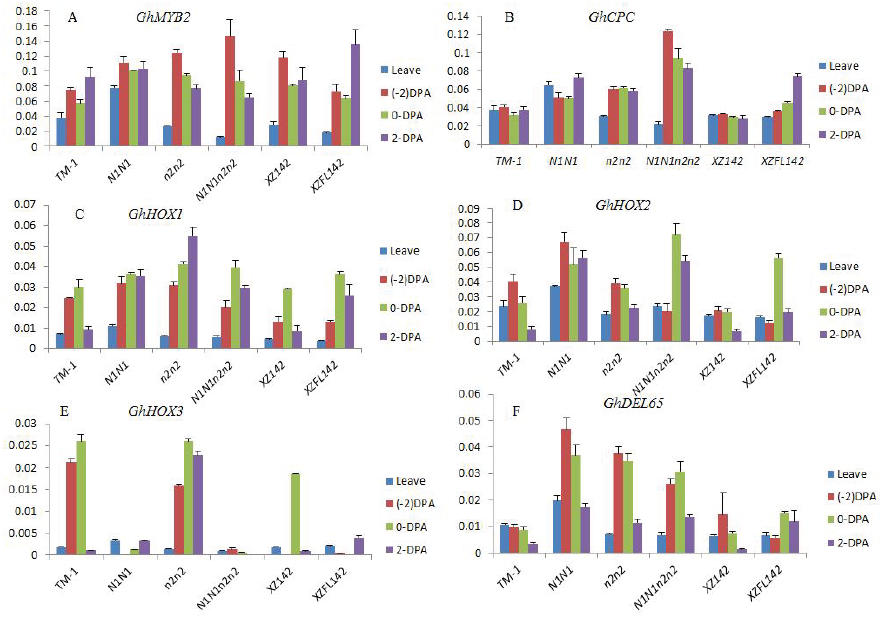
Transcripts profiling for potential fiber initiation related genes. (A) Gene expression for *GhMYB2* (*26*) in different varieties and development stages. (B) Gene expression for *GhCPC* (*S3*) in different varieties and development stages. (C) Gene expression for *GhHOX1(26)* in different varieties and development stages. (D) Gene expression for *GhHOX2* (*26*) in different varieties and development stages. (E) Gene expression for *GhHOX3 (26)* in different varieties and development stages. (F) Gene expression for *GhDEL65* (*S4*) in different varieties and development stages.

**Fig. S3.**
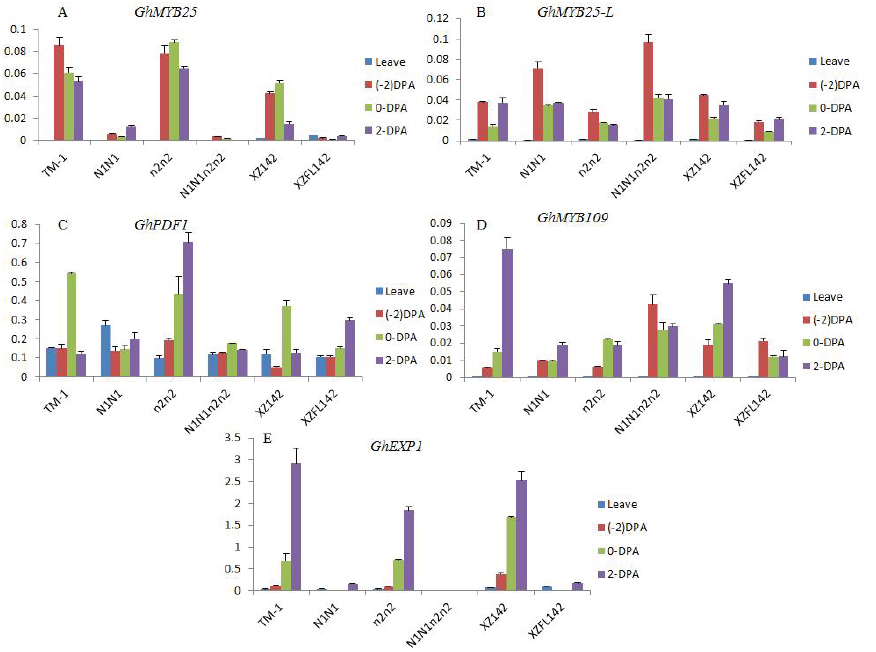
Transcripts profiling for potential fiber elongation related genes. (A) Gene expression for *GhMYB25* (*29*) in different varieties and development stages. (B) Gene expression for *GhMYB25-L* (*29*) in different varieties and development stages. (C) Gene expression for *GhPDF1* (*S5*) in different varieties and development stages. (D) Gene expression for *GhMYB109* *(S6)* in different varieties and development stages. (E) Gene expression for *GhEXP1* (*S7*) in different varieties and development stages.

**Table S1.**
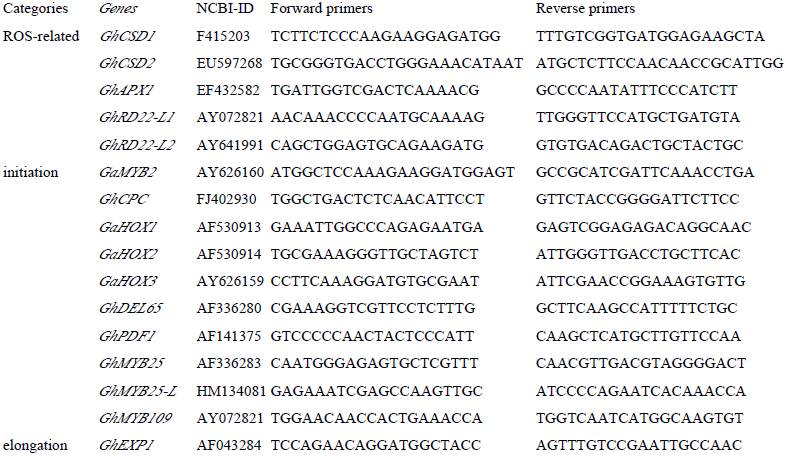
QRT-PCR primers for potential fiber initiation and development genes.

